# Adaptive metabolic reprogramming of the brain tissue during late-stage *Trypanosome brucei* infection maintains host learning and memory

**DOI:** 10.64898/2026.05.15.725345

**Authors:** Nadin Fathallah, Chloe Barnes, Rebecca Chatwin, Jayde Whittingham-Dowd, John Worthington, Lucy Jackson-Jones, Neil Dawson, Michael D. Urbaniak

## Abstract

Human African Trypanosomiasis (HAT) is a two-stage infection caused by *Trypanosoma brucei* ssp. In stage I the trypanosomes are in blood, lymph and tissue interstitial space and the infection progresses to stage II when the parasites enter the central nervous system (CNS), resulting in behavioural aberrations that proceed to coma and death. Here, we use a bioluminescent murine model of HAT to examine parasite localisation and the changes in host brain gene expression, metabolism and function, and behaviour that occur over the course of the infection. The murine HAT model reproduces the decrease in brain tryptophan seen in clinical samples, and we report for the first time an unprecedented 1.8-fold decrease in global brain glucose metabolism in stage II infection. These metabolic changes are accompanied by an 18-fold decrease in brain insulin transcripts without changes in pathways regulating the cellular responses to insulin. By contrast, genes involved in fatty acid and lipid metabolism are upregulated in the brain during stage II infection. Moreover, we show that transcriptional programmes regulating mitochondrial metabolism dynamically adapts across the time course of HAT infection, ultimately leading to a transcriptional programme that diverts host brain metabolism away from glycolysis during stage II infection. Overall, our data demonstrate a reprogramming of brain energy metabolism during stage II HAT infection that favours the utilization of fatty acids and lipids to meet the energy demands of the brain, with a reduced reliance on glucose metabolism. Despite the profound neurometabolic changes observed, host anxiety-like behaviour is unchanged and episodic learning and memory is not impaired, suggesting that brain metabolic reprogramming enables the utilisation of adipose reserves to maintain core brain functions. These finding may explain the progressive onset of neurological symptoms in HAT patients and inform the development therapeutic interventions to alleviate them.

**Author Summary:** Human African Trypanosomiasis is classically characterised as being a two-stage infection, stage I where the extracellular *Trypanosoma brucei* multiply in the blood, lymph and peripheral tissues, and stage II where parasite cross the blood-brain barrier (BBB) causing neurological symptoms and eventually death. Using a murine model of HAT we show that in stage II the key brain metabolite tryptophan is depleted and cerebral glucose utilisation is decreased, accompanied by extensive metabolic transcriptome reprogramming of the cerebral tissue during stage II infection. Despite this we see no significant change in mouse anxiety-like behaviour or learning and memory. Our data are consistent with the brain switching from glucose as the primary energy source, instead utilising the products of lipolysis to maintain essential brain functions. This new understanding of the neurometabolic changes that occur in stage II HAT may help to develop new treatments for the neurological symptoms that affect patients at this stage of the disease.

## Introduction

The African trypanosomes *Trypanosoma brucei gambiense* and *Trypanosoma brucei rhodesiense* are infective to humans causing the neglected disease Human African Trypanosomiasis (HAT), also known as African Sleeping Sickness. *T. brucei ssp.* are transmitted in sub-Saharan Africa by the bite of infected tsetse fly (*Glossina ssp.*), with an estimated 65 million people at risk in the tsetse belt. In the first stage of HAT the parasite multiplies in the blood, lymph and interstitial space, and in the second stage the parasite enters the central nervous system (CNS) causing severe neurological disturbances and, if untreated, death [1]. As trypanosomes evade the host immune system by regularly switching their surface coat, an effective vaccine is yet to be developed. Despite this, over the past two decades the number of cases of HAT has decreased by >95% due to increased vector control, disease surveillance, and the development of improved therapeutics [2].

In stage I (S1) HAT *T. brucei* occupy diverse tissues within the host resulting in fever, weight loss, anaemia, headache and fatigue. Localisation to numerous tissues has been observed in HAT animal models, by histopathology and the use of bioluminescent parasites [3, 4]. The skin is the initial point of entry of tsetse transmitted infections, and skin localisation is critical for onward parasite transmission and represents an important anatomical reservoir [5, 6]. A high parasite burden has also been identified in adipose tissues in mouse models, with adipose-resident parasites adapting their metabolism to increase sterol and lipid metabolism and β-oxidation, allowing the parasites to utilise the fatty acids available in the local tissue as a source of energy [7]. Loss of adipose tissue is a major contributor to weight loss in trypanosome infection, with higher adipocyte lipolysis promoting host survival [8].

In stage II (S2) HAT the parasite crosses the blood brain barrier (BBB) and invades the CNS resulting in neurological disruption including psychiatric symptoms and a characteristic disturbance of sleeping patterns, which gave rise to the name African Sleeping Sickness. Despite the reported low parasite burden in the brain as compared to blood and other tissues [7] the most severe pathology is always associated with brain colonisation [1]. However, the basis of the neurological symptoms observed is not well understood. Importantly, neurological impairments persist after infection, with patients successfully cured of HAT experiencing long-term sleep disturbance, ataxia and psychiatric disorders [9]. Improved understanding of the connection between infection and the resultant neuropsychopathology may enable the identification of new ways to alleviate these symptoms to improve therapeutic outcomes.

*T. brucei* are auxotrophs that obtain essential nutrients including glucose and amino acids from their hosts. *In vivo* infection models of *T. b gambiense* show increased tryptophan turnover in the brain [10], and metabolomics studies of S2 HAT patient samples confirm that a significant decrease in tryptophan levels occurs in the cerebrospinal fluid (CSF) [11]. *T. brucei* cultured *in vitro* significantly deplete tryptophan from the media [12] which they use in both protein synthesis and transamination [13], secreting the unused biochemical intermediates of the transamination reactions as aromatic keto-acids [10, 14, 15]. These secreted metabolites significantly impact host physiology. For example, the trypanosome-derived tryptophan metabolite 3-indole pyruvate modulates host immunity [14, 15], whilst its metabolite tryptophol has sleep-inducing properties [10]. Tryptophan is the precursor of the kynurenine pathway which has both neuroprotective and neurotoxic branches that impact HAT pathology [16]. Tryptophan is also the precursor of the neurotransmitter serotonin and the hormone melatonin that regulate host behaviour, sleep [17] and immune responses in the brain [18]. Thus, tryptophan metabolism by trypanosomes in the CNS may directly interfere with the host immune response and neurotransmission, through both the depletion of tryptophan and the secretion of neuroactive and immune-modulatory tryptophan-derived metabolites.

Here, we show that an established bioluminescent murine model of S1 and S2 HAT [19] reproduces the changes in tryptophan metabolism observed in clinical samples, and examine how changes in tryptophan metabolism relate to host brain activity and behaviour. Surprisingly, while metabolic brain imaging revealed a significant global reduction in brain glucose utilisation, no significant effect on anxiety-like behaviour or episodic memory was found. Our transcriptional analysis demonstrated an increase in host gene expression in fatty acid and lipid metabolism pathways, consistent with a transcriptional switch away from glycolysis during S2 infection towards the use of lipids to meet the energy demands of the brain. This new understanding of infection-induced brain metabolic reprogramming may lead to improved therapeutic outcomes for the neuropsychopathology seen in HAT.

## Results

### Measuring changes in metabolism and behaviour in a bioluminescent murine model of HAT

To assess the effect of *T. brucei* infection on host metabolism and behaviour we use an established experimental murine HAT model infected with a pleomorphic *T. brucei* VSL2 strain that highly expresses a bioluminescent, red-shifted luciferase [19]. In this model, an intraperitoneal (*i.p.*) injection results in a S1 infection from day 0-7, after which the parasites progressively cross the BBB and establish an early-stage II (ES2) infection between day 7-14, with a late-stage II (LS2) infection occurring after day 14 and a survival time of 25-35 days post infection. Tissues were collected from control and infected mice at S1, ES2 and LS2 for *ex vivo* imaging to characterise parasite tissue localisation and to obtain material for metabolomic and transcriptomic analysis, with behaviour testing conducted at LS2 (Supplementary Figure S1).

### *T. brucei* are widely distributed in host tissues during infection

*Ex vivo* imaging of diverse tissues revealed a widespread parasite distribution that varied between tissues (Figure 1A-C, Supplementary Figures S2-5). The total parasite burden increased from early to late infection, reflecting the increased parasite load that occurs after the first peak in parasitaemia (Figure 1C). Some inter-animal variation in parasite burden and distribution was observed at S1, while burdens became more consistent in ES2 and LS2. We used mean bioluminescence intensity (I_m_) to classify tissues as being of low (Log_10_ I_m_ < 5.5), medium (5.5 < Log_10_ I_m_ < 7.0), or high (Log_10_ I_m_ > 7.0) parasite burden. At S1 a consistently high parasite burden occurred only in the gut draining mesenteric lymph node (mLN), with medium parasite burden observed in adipose tissue, colon, small intestine (SI) and spleen, and low but detectable parasite burden found in other tissues. As the infection progressed the number of tissues with a medium or high parasite burden increased, with high parasite burden found in adipose, colon, mLN, muscle, SI and spleen during S2. Bioluminescence intensity within the brain was significantly lower (p < 0.05) than that in other tissues (Figure 1B and C).

**Figure 1.**
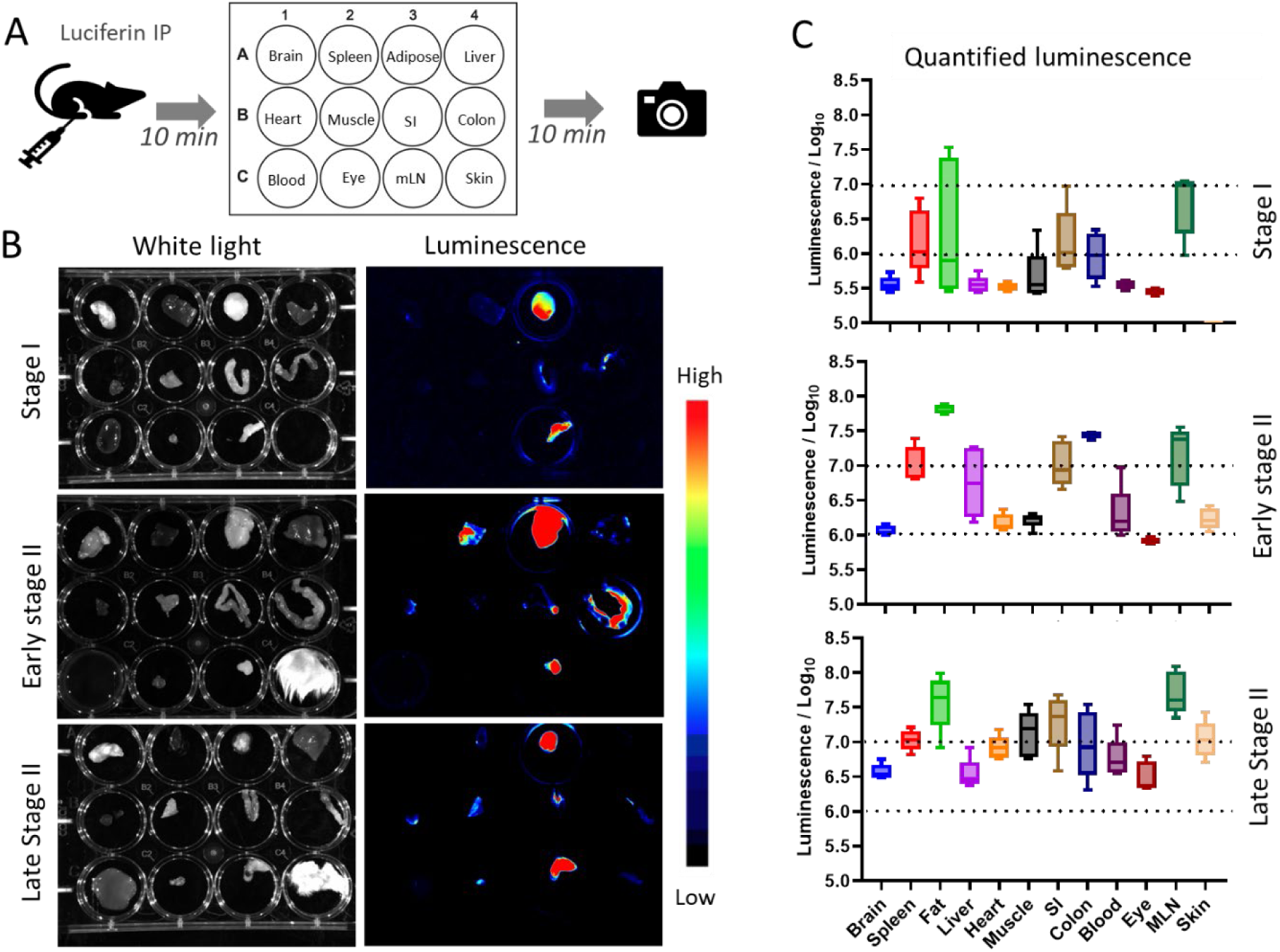
*Ex vivo* imaging of bioluminescent *T. brucei* VSL2 in mouse tissues. **A.** Mice were injected *i.p.* with Luciferin at stage I (day 7), early stage II (day 14) and late stage II (day 22) of infection prior to humane sacrifice and *ex vivo* imaging in 12 well plates in a standard layout; SI – small intestine, LI – large intestine, mLN – mesenteric lymph node. **B.** White light and luminescence imaging of tissue in plate format; luminescence visualised as heat-spectrum, typical examples shown. **C.** Quantification of average luminescence intensity showing interquartile range (box), range (whiskers) and mean (X). n = 5. Skin samples were omitted from day 7 imaging.

### Brain tryptophan levels show a biphasic change during infection

Given the reports of significant decreases in tryptophan in S2 HAT patients [11], we aimed to determine the effect of *T. brucei* infection on brain tryptophan levels in our murine HAT model using targeted metabolomics. Brain samples of infected and control animals at S1, ES2 and LS2 of infection (Figure 2A) were analysed by high-performance liquid chromatography (HPLC) using fluorescence detection of tryptophan [20]. The data reveals a biphasic change in brain tryptophan levels in the infected groups relative to controls (Figure 2B), with the level of tryptophan significantly *increased* during S1 (2.1-fold, p < 0.0001) and *decreased* during ES2 and LS2 (1.8-fold, p < 0.01) infection. Brain samples from ES2 and LS2 of infection were also analysed by liquid chromatography–mass spectrometry (LC-MS/MS) to directly quantify the abundance of tryptophan [21] and confirm the specificity of the fluorometric HPLC analysis. The ES2 brain samples showed a similar but non-significant decrease (1.4-fold, p > 0.05) in tryptophan, with a significant decrease (1.7-fold, p < 0.05) observed in LS2 samples as compared to control animals (Figure 2C), confirming the HPLC data by an orthogonal technique.

**Figure 2.**
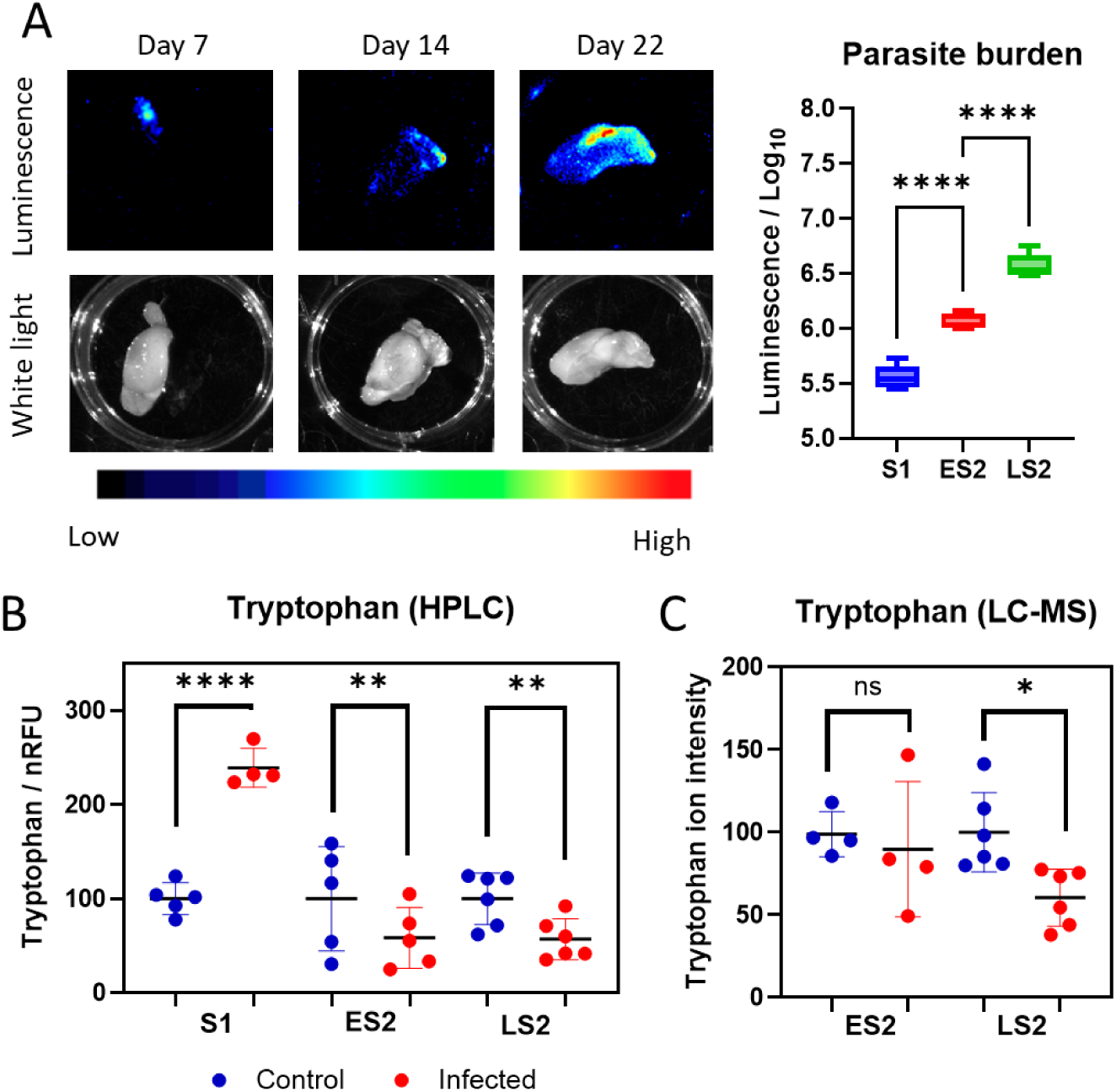
Brain tryptophan levels show a biphasic change over the course of the infection. **A**. *Ex vivo* imaging of bioluminescent *T. brucei* VSL2 in mouse CD-1 brain tissues with quantification of average luminescence intensity (details as Figure 2). **B.** HPLC analysis of brain tryptophan levels; S1 – stage I, day 7, n = 4, ES2 – early-stage II day 14, n = 5; LS2 – late-stage II, day 22, n = 6. **C.** LC-MS analysis of Stage II brain samples; ES2 – early-stage II day 14, n = 4; LS2 late-stage II, day 22, n = 6. ns p > 0.05, * p < 0.05, ** p < 0.01, *** p < 0.001 and **** p < 0.0001 significant difference between control and infected mice.

### Tryptophan levels in peripheral infected tissues are not associated with parasite burden

To determine whether tryptophan depletion is a brain specific or a multi-organ feature of trypanosome infection, analysis of additional tissues that display different parasite burdens at S1 and LS2 was undertaken (Figure 3A). The liver is the predominant source of tryptophan 2,3-dioxygenase (TDO), the first enzyme initiating the kynurenine pathway which accounts for ∼95% of tryptophan metabolism in humans. A 1.9-fold *increase* in liver tryptophan levels was observed at both S1 (p < 0.001) and LS2 (p < 0.0001), despite the increase from low to medium parasite burden across these stages (Figure 3A). By contrast, a biphasic tryptophan response was seen in the muscle and spleen across the infection time course. In muscle a no-significant change in tryptophan levels was observed in S1, contrasting with a pronounced 2.3-fold *increase* (p < 0.0001) at high parasite burden in LS2 (Figure 3B). In the spleen a significant 1.3-fold *decrease* (p < 0.05) in tryptophan levels was observed in S1 at medium parasite burden, which altered to a 1.3-fold *increase* (p < 0.01) at high parasite burden in LS2 (Figure 3C). These data show that the impact of *T. brucei* infection on tryptophan levels across the time course of infection is tissue specific. Moreover, the increased tryptophan levels observed in the liver, muscle and spleen during LS2 do not align with decreased tryptophan observed in the brain at this stage, suggesting that brain tryptophan levels are largely uncoupled from peripheral tryptophan availability during LS2 of infection. In addition, the changes in tryptophan levels seen in the brain, spleen, muscle and liver induced by *T. brucei* infection do not show a consistent association with tissue parasite burden, with the brain being notable as the only tissue where a decreased tryptophan levels occurs in LS2 of infection.

**Figure 3.**
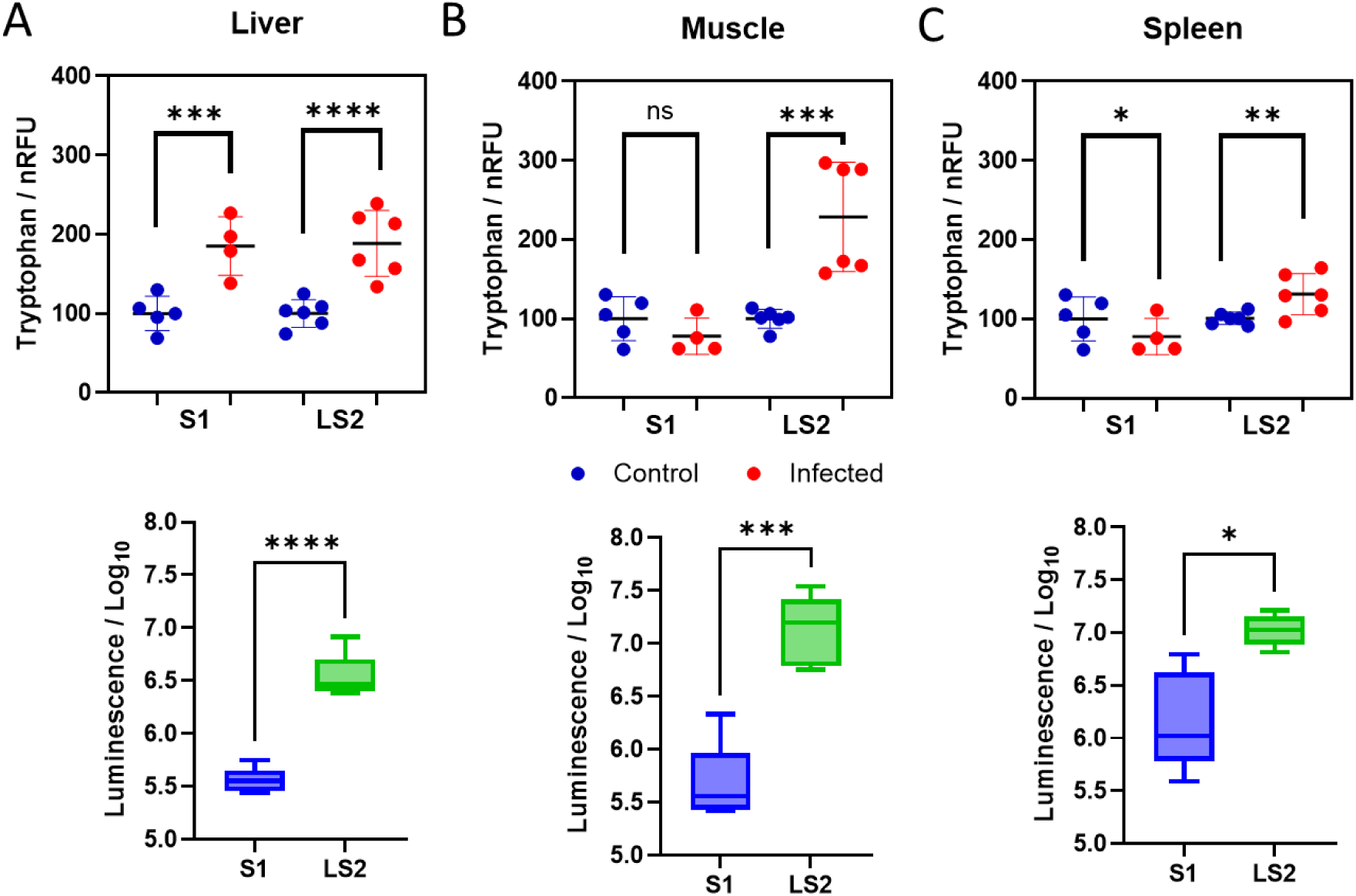
Tissue tryptophan levels are altered over the course of the *T. brucei* infection. **A.** Liver**, B.** Muscle**, C.** Spleen. Upper panel – HPLC analysis of tryptophan abundance; S1 – stage I, day 7, control n = 5, infected n = 4; LS2 – late stage II, day 22, n = 6. Lower panel – Quantification of *T. brucei* VSL2 luminescence in tissues (details as Figure 2), n = 6. ns p > 0.01, * p < 0.05, ** p < 0.01, *** p < 0.001 and **** p < 0.0001 significant difference between control and infected mice.

### Global cerebral glucose utilisation is reduced in late stage II infection

To assess the effect of brain parasite localisation and the altered tryptophan levels seen during LS2 infection on brain function we performed brain imaging to measure cerebral glucose metabolism [22]. Brain imaging using ^14^C 2-deoxy-glucose (2-DG) provides a quantitative measure of brain tissue glucose utilisation, identifying metabolically active regions of cerebral tissue that reflects regional neuronal activity.

Surprisingly, we found evidence for an unprecedented widespread reduction in cerebral glucose utilisation in the brains of S2 *T. brucei* infected mice across the whole brain (whole brain average: 1.8-fold decrease, p < 0.001) (Figure 4A) and evident across multiple regions of diverse neural systems (Figure 4B-F and I, Supplemental Table S1). This included reduced glucose utilisation in regions of the prefrontal cortex, basal ganglia, hippocampus and in the serotonergic dorsal raphé. *T. brucei* rapidly consumes glucose and can take up 2-DG [23] and therefore parasite presence in the cerebral tissue could increase the apparent rate of glucose utilisation within the infected brains. However, this is contrary to our observation of decreased 2-DG utilisation. High plasma glucose concentrations can theoretically outcompete transport of ^14^C-2-DG across the BBB, which could result in a global reduction in brain ^14^C-2-DG levels [24]. However, as we found *T. brucei* infection induced a small but significant *reduction* (1.2-fold, p < 0.05) in circulating plasma glucose levels (Figure 4G), this suggests our brain imaging results are valid. Moreover, plasma ^14^C-2-DG levels were not significantly different between the experimental groups (Figure 4H), further supporting the validity of our observation that S2 *T. brucei* infection significantly decreases the use of glucose by cerebral tissue.

**Figure 4.**
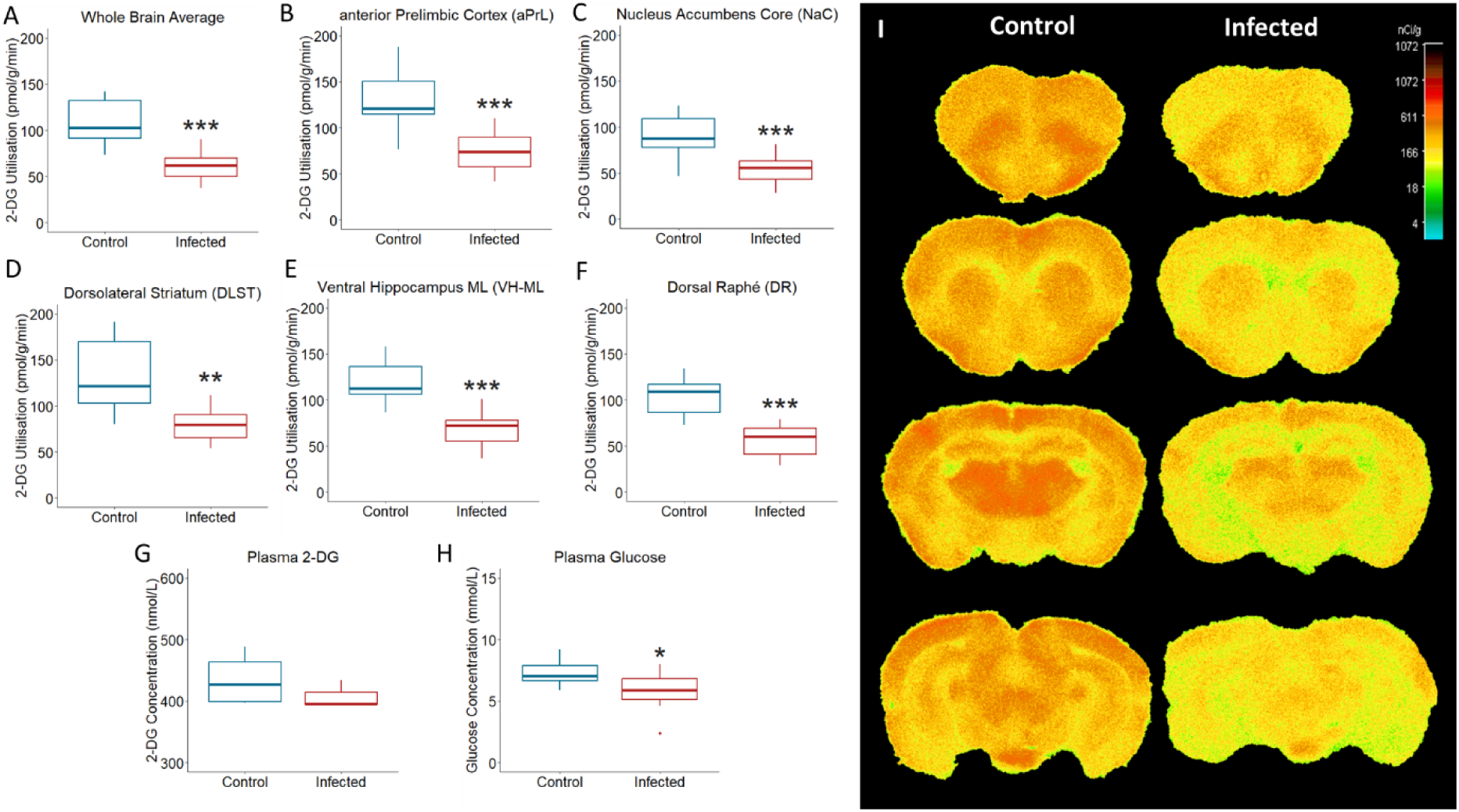
Late Stage II *T. brucei* infection induces a widespread reduction in glucose metabolism within the brain. Glucose utilisation (pmol/g/min), measured using ^14^C-2-deoxyglucose (^14^C-2-DG) autoradiographic metabolic imaging in *T. brucei* infected mice in LS2 (day 22, infected n = 13, control n = 12). **A**. whole brain; **B-F** broad range of brain regions. **G.** Plasma glucose levels (mmol/L) in *T. brucei* infected mice in LS2. **H.** Plasma ^14^C-2-deoxyglucose concentrations. I. Representative, false-colour brain images of ^14^C-2-DG levels (nCi/g) across the brain. *p < 0.05, **p < 0.01 and *** p <0.001 significant difference from control mice.

### Host episodic learning and memory are not modified in stage II infection

To determine the impact of *T. brucei* infection on host behaviour in our murine HAT model we performed behavioural tests during LS2 infection, as this is when neurological symptoms occur in HAT patients. The open field test (OFT) was employed to assess exploratory locomotor and anxiety-like behaviour. We found significant evidence for reduced locomotor activity in infected animals in LS2 compared to control animals (Figure 5A). However, we found no evidence for altered anxiety-like behaviour in infected animals, as indicated by no significant change in the amount of time spent in the centre zone of the OFT arena (Figure 5B).

**Figure 5.**
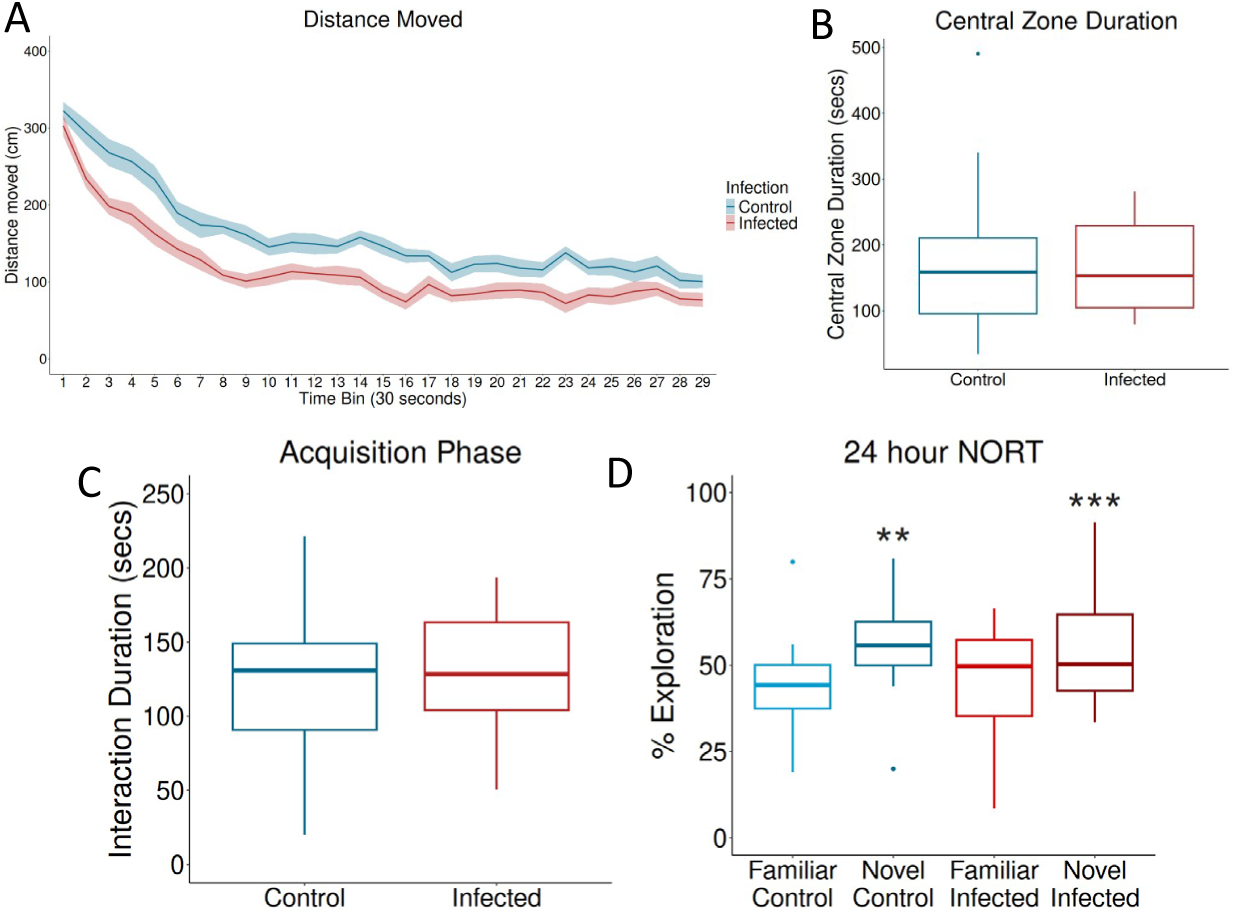
Host episodic learning and memory, and anxiety-like behaviour, are not altered during stage II *T. brucei* infection. **A.** Locomotor activity is significantly (p < 0.001, repeated measures ANOVA) reduced in LS2 infected animals (day 20, n = 6) as compared to control animals in the Open Field Test (OFT). Data are shown as the mean ± SEM per 30 second time bin. **B.** Anxiety-like behaviour, as reflected by the total duration (seconds) spent in the central zone during the OFT, is not significantly altered in LS2 infected animals (day 20, n = 6). **C.** Novel object recognition test (NORT): The amount of time spent exploring the objects is not different between LS2 infected and non-infected mice during the acquisition phase (day 21, n = 6). **D.** Both LS2 infected (day 22, n= 6) and control animals show object recognition memory 24 hours after exposure to the objects during the acquisition phase, as evidenced by spending significantly more time exploring the novel as compared to the familiar object, ** p < 0.01, ***p < 0.001 significant difference in the percentage exploration for novel as compared to the familiar object.

**Figure 6.**
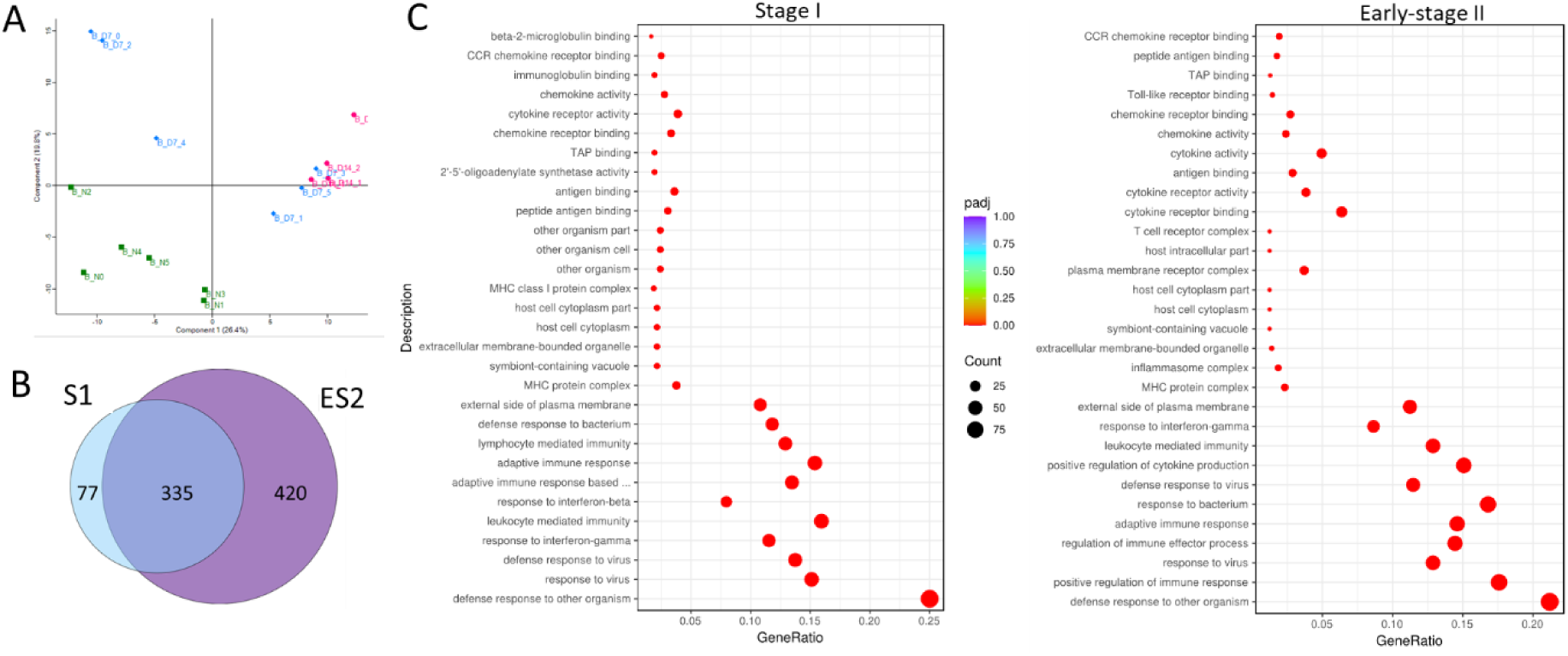
RNAseq analysis of changes in the host brain transcriptome during infection. RNAseq data for right hemisphere of brains from naive (n = 6) and *T. brucei* infected animals at S1 (day 7, n = 6) and ES2 (day 14, n = 4). **A.** Principal Component Analysis (PCA) demonstrate distinct clustering of naïve and infected cohorts. **B.** Venn of Differentially Expressed Genes (DEGs) between naïve and infected cohorts. **C.** GO term enrichment analysis dot plot comparing naive to infected samples at S1 and ES2.

Next, we sought to characterise the impact of infection on episodic learning and memory using the novel object recognition test (NORT). During the acquisition phase, where animals explore the objects, we found no evidence for a difference in the amount of time spent exploring the objects between control and infected animals (Figure 5C), supporting no impact on task engagement. 24 hours after familiarisation we tested object recognition memory. Both control and infected animals spent significantly more time exploring the novel as compared to the familiar object at this time (Figure 5D), indicating that animals from both groups had formed a memory of the familiar object. Overall, this suggests that LS2 *T. brucei* infection did not result in significant episodic learning and memory deficits, despite its impact on brain glucose metabolism in areas involved in learning and memory.

### RNAseq analysis of brain tissue reveals host metabolic reprogramming across the time course of *T. b. brucei* infection

To assess the effect of *T. brucei* infection on host gene expression in the brain we performed RNAseq analysis of control and infected brain tissue during S1 and ES2 of infection (Supplementary Table S2). The biological replicates of each stage of the infection had strong correlation (Pearson correlation 0.94 – 0.98), and Principal Component Analysis (PCA) showed that naive animals clustered separately from infected animals (Figure 7A). However, the distinction between some S1 and ES2 animals was less pronounced, which may reflect natural variability in the time at which the parasites cross the BBB and some of the overlapping changes in gene expression seen in infected animals at both time points. Analysis of the Differentially Expressed Genes (DEGs) between naïve and infected cohorts revealed that the DEGs in S1 (412) and ES2 (755) were predominately upregulated in infected cohorts (Supplementary Table S3 & S4). A predominance of upregulated DEGs in brain tissue in response to *T. brucei* infection was previously observed by Quintana *et al* [25]. In S1 there were 401 significantly upregulated and 11 significantly downregulated transcripts, and ES2 there were 721 significantly upregulated and 34 significantly downregulated transcripts (Log_2_(Fold-Change) ≥ |1| and p < 0.05) in comparison to naïve animals. The majority of DEGs identified in S1 were also detected at ES2 (355/412) whilst an additional 420 were specific to ES2 infection (Figure 7B) providing evidence that the host response alters between S1 and ES2.

**Figure 7.**
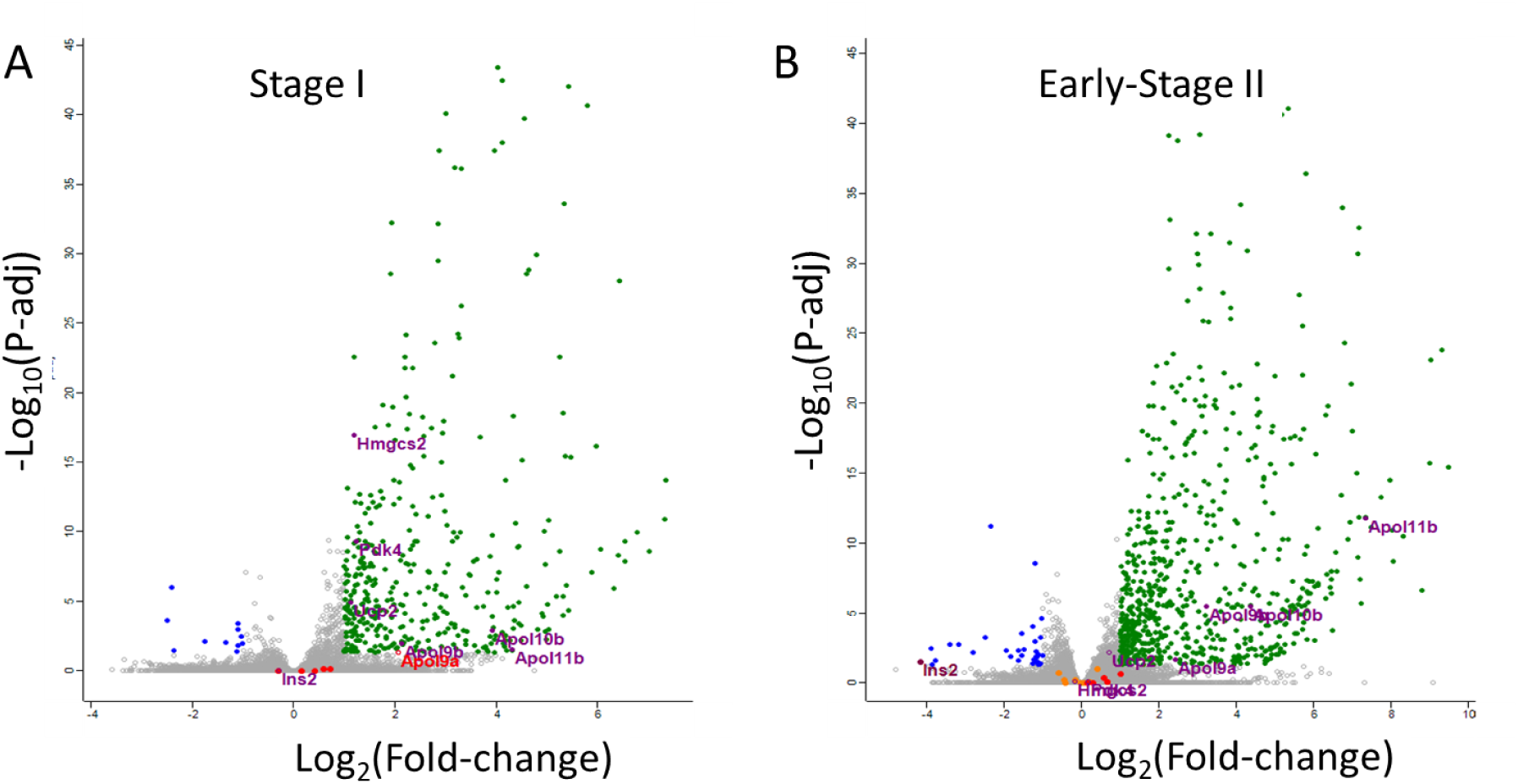
Volcano plots of host brain RNAseq data showing differentially expressed genes (DEGs) in stage I (S1) and early-stage II (ES2) mice. RNA sequencing of mouse brains (left hemisphere) comparing naïve (n = 6) with S1 (day 7, n = 6), ES2 (day 14, n = 4) infections. Green – Upregulated DEGs; Blue – Downregulated DEGs; Orange – Glucose transporters *Slc2a1* – *Slc2a13*; Red – Tryptophan metabolising enzymes *Ido1*, *Ido2*, *Tdo2*, *Tph1* and *Tph2;* Purple – Insulin II - *Ins2,* apolipoprotein L9a - *ApoL9a*, apolipoprotein L9b - *ApoL9b*, apolipoprotein L10b - *ApoL10b,* apolipoprotein L11b - *ApoL11b* uncoupling protein 2 - *Upc2,* pyruvate dehydrogenase kinase 4 - *Pdk4,* 3-hydroxy-3-methylglutaryl-CoA synthase 2 - *Hmgcs2,* lipolysis-stimulated lipoprotein receptor –*Lsr*.

As expected, Gene Ontology (GO) term enrichment analysis of DEGs between naïve and infected samples (Figure 7C) identified GO terms predominantly related to the immune response against infection in S1 that became more widespread in ES2 [25], in-line with the increased parasite burden in the brain. In the brains of S1 animals (versus naïve) these enriched GO pathways included those linked to activation of the innate (GO:0019882) and adaptive (GO:0002250) immune response, positive regulation of cytokine production (GO:000819) and immune response (GO:0050778), and response to interferon gamma (GO:0034341) and beta (GO:0035456). The dominance of immune responses in S1 of the infection may be due to the combination of an expansion of brain resident immune cell populations and the recruitment of peripheral immune cell populations to the brain [26]. Upregulated expression of genes in these immune GO pathways was further enhanced in ES2 versus S1 animals. These larger changes in ES2, which is also known as the meningoencephalitis stage, likely represent direct neuroinflammation caused by parasite infiltration into the brain at this stage and degradation of the BBB [25, 26].

Amongst the small number of genes significantly downregulated in the infected brains in comparison to non-infected controls there were only 5 out of 11 in S1 with a defined function compared to 24 out of 43 in ES2. There was no evidence to suggest that glucose transporter gene expression (*Slc2a1* – *Slc2a5*) was significantly altered in ES2, suggesting that glucose transport in the CNS was not impaired by infection. However, consistent with the reduced cerebral glucose metabolism we identified in LS2 mice, expression levels of the Insulin II (*Ins2*) gene, which regulates cerebral tissue glucose use [27], was dramatically decreased by 18-fold in ES2 brains (Figure 7). By contrast, the expression of genes in the GO cellular response pathways to insulin signalling (GO:1900076) and the regulation of glucose import in response to insulin stimulus (GO:2001273) were not altered in the brains of ES2 mice, suggesting that decreased glucose use in the brain was driven by attenuated brain Insulin II availability, rather than through a lack of tissue insulin sensitivity.

Intriguingly, in both stages of infection, the expression of mitochondrial encoded genes was not significantly altered in comparison to naïve animals, suggesting that mitochondria number were not significantly altered in the brain tissue. However, the expression of several nuclear encoded mitochondrial genes known to regulate mitochondrial metabolic function were significantly increased in S1 brains. This included a 2.4-fold increase in *Pyruvate dehydrogenase kinase 4* (*Pdk4*) and 2.4-fold increase in *3-hydroxy-3-methylglutaryl-CoA synthase 2* (*Hmgcs2*) expression in S1 animals. Comparison of ES2 versus S1 brains revealed further evidence of metabolic reprogramming of the brain tissue across this time course, evidenced by significantly decreased expression of both *Pdk*4 (2.1-fold reduction) and *Hmgcs2* (2.6-fold reduction) during ES2 infection, returning these to similar levels of expression as seen in naïve animals. By contrast, expression of the mitochondrial glucose metabolism regulating gene *Uncoupling protein 2* (*Ucp2*), which uncouples mitochondrial ATP production from glycolysis and promotes the use of alternative substrates including fatty acid and glutamine, was significantly increased in the brains of both S1 (2.2-fold increase) and ES2 (1.7-fold increase) animals. Overall, these data support dynamic, adaptive regulation of mitochondrial metabolism across the infection time course, that aligns with the reduced cerebral glucose metabolism we have identified through metabolic brain imaging in LS2 animals.

The reduced level of brain insulin (*Ins2*) is consistent with the reduced glucose utilisation we observe in LS2 brains, and will promote the metabolic use of lipids through increased lipolysis and release of fatty acids [28]. Interestingly, we also observe a large and significant increase at ES2 in *Apolipoprotein L11b* (*ApoL11b* – 162-fold) and *Apolipoprotein 10b* (*ApoL10b* – 21-fold) expression, both involved in the formation of high-density lipoproteins for lipid transport, and a corresponding increase in the *Lipolysis-stimulated lipoprotein receptor* (*Lsr* – 2.1-fold), a membrane protein that promotes cellular uptake of triglyceride-rich lipoproteins. Intriguingly, the human homologue *APOL1* is a trypanosome lytic factor that is lethal to all African trypanosomes except for the two human infective sub-species that cause HAT [29], which suggests these genes may be upregulated in an attempt to clear the brain of the parasites. Interestingly, there is also a significant increase in the *Apolipoprotein L9* isoforms *ApoL9a* and *ApoL9b* in ES2 mouse brains, which are interferon (IFN) stimulated genes that are upregulated in neurons in response to local production of type I IFN [30]. Similarly increased expression of these genes were also observed in trypanosome infected adipose tissue by Machado *et al* [8], further supporting a link between increased lipolysis and the immune response during infection. Interestingly, transcript levels of genes for the tryptophan metabolising enzymes, *Ido1*, *Ido2*, *Tdo2*, *Tph1* and *Tph2*, were not significantly altered in either S1 or ES2 brains, suggesting that alterations in host brain metabolism of tryptophan did not underlie the differences seen in the brain tissue levels of tryptophan seen during infection.

### Transcriptomic changes in brain-resident *T. brucei* also support parasite metabolic adaption

To examine changes in the parasite transcriptome over the course of infection the same RNAseq data from infected brain tissue were aligned to the 35 Mb *T. brucei* genome. A low percentage of reads (0.03 – 0.18%) from both S1 and ES2 samples aligned with the *T. brucei* genome, identifying 6,378 transcripts (Supplementary Table S5) from the 10,239 annotated genes with a moderate but significant correlation between biological replicates (Pearson correlation 0.62 – 0.84), and no clear distinction between the parasites at the different stages of infection based on PCA (Figure 8A), suggesting that gene transcription was largely conserved in *T. brucei* across S1 and ES2. However, analysis of *T. brucei* DEGs between S1 and ES2 cohorts revealed 26 significantly down-regulated genes and 72 significantly up-regulated genes (Log_2_(Fold-Change) ≥ |1| and p < 0.05) (Figure 8B, Supplementary Table S6). This data confirms the presence of transcriptionally active parasites within the brain tissue at both S1 and ES2 as observed by bioluminescence (Figure 2), and that the transcriptome of brain-resident parasites changes across the time course of infection.

**Figure 8.**
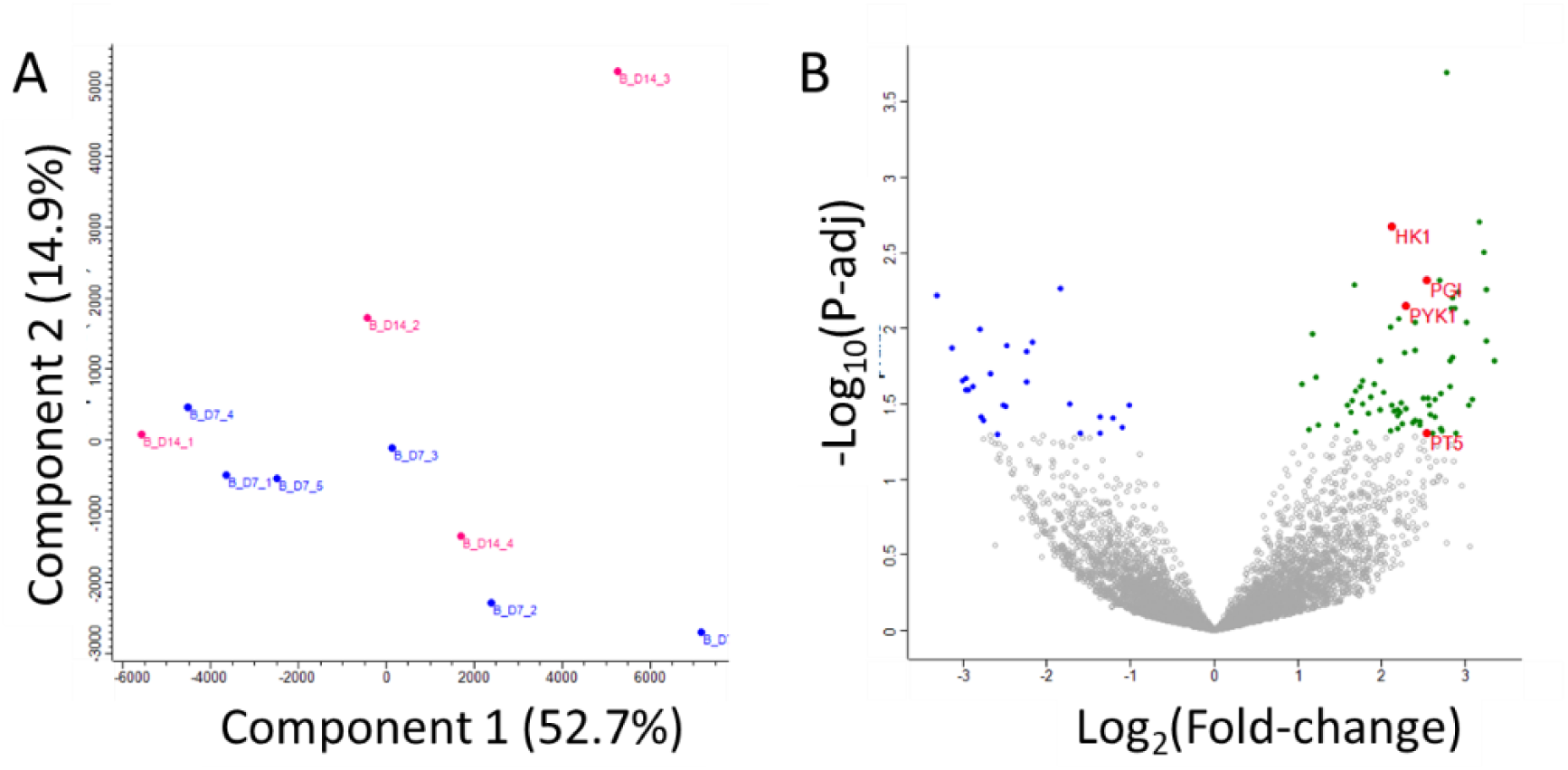
RNAseq analysis of changes in the *T. brucei* transcriptome during infection. RNA sequencing of infected mouse brains at S1 (day 7, n = 6) and ES2 (day 14, n = 4) aligned to the *T. brucei* genome. **A.** Principal Component Analysis (PCA) shows no distinct clustering of S1 and ES2 biological replicates. **B.** Volcano plot of differentially expressed genes (DEGs), Log_2_ Fold-Change ≥ 1 and p > 0.05. Green – Upregulated DEGs; Blue – Downregulated DEGs; Red – *hexokinase 1* (*HK1*), *phosphoglucose isomerase* (*PGI)*, *pyruvate kinase 1* (*PYK1)*, *mitochondrial malate kinase* (*MEM), pyruvate transporter* (*PT5)*.

Gene Ontology (GO) term enrichment analysis of *T. brucei* DEGs between ES2 versus S1 (Supplemental Table S8 & S9) identified upregulated GO terms for glycolytic processes (GO:006096) and pyruvate metabolic processes (GO:006090) with genes included *hexokinase 1* (*HK1)*, *phosphoglucose isomerase* (*PGI*), *pyruvate kinase 1* (*PYK1)*, *mitochondrial malate kinase* (*MEM)* and the *pyruvate transporter* (*PT5),* which mediates pyruvate efflux, in ES2 trypanosomes (Figure 8B). No significant change was observed for the key tryptophan transamination enzyme *cytosolic aspartate aminotransferase* (*cASAT*) [14] between S1 and ES2. The increase in parasite enzymes critical for energy metabolism in ES2 compared to S1 is consistent with adaption to nutritional stress caused by host metabolic reprogramming and reduced glucose availability (as supported by reduced plasma glucose availability in LS2 infected animals), and points to increased pyruvate production and efflux from the parasites.

## Discussion

Tryptophan and its metabolites mediate host immune response to HAT [14, 15] and influence pathology [10, 16]. We have shown that an experimental murine HAT model using the pleomorphic bioluminescent *T. brucei* VSL2 strain [19] reproduces the perturbation of brain tryptophan observed previously in HAT animal models and patients [10, 11]. Brain tryptophan levels increased in S1 of infection before decreasing in ES2 and LS2 of infection. This biphasic trend is in agreement with LC-MS/MS metabolomic analysis of clinical HAT CSF samples by Vincent *et al.* [11], where decreased tryptophan was observed during S2. While Vincent *et al.* also observed a modest increase in CSF tryptophan during S1 HAT it did not reach statistical significance, potentially reflecting the greater variability in time since infection in their clinical samples as compared to our time-matched murine model.

The perturbation of brain tryptophan is decoupled from tryptophan levels within peripheral tissues, as the liver, muscle and spleen all show a significant increase in tryptophan in LS2, which is opposite to that observed in the brain. Interestingly, alterations in tryptophan in both the brain and peripheral tissue did not correspond with tissue parasite burden, despite evidence that *T. brucei* significantly deplete tryptophan from cell culture media [12]. Our RNAseq data suggests that alterations in host brain tryptophan metabolism are also unlikely to account for the observed changes in brain tryptophan, as the transcript levels of tryptophan metabolising enzymes were not significantly altered during the infection. The driving force for the clinically observed decreased brain tryptophan levels in S2 HAT thus remains unclear and warrants further investigation.

The healthy mammalian brain is responsible for ∼25% of the total consumption of glucose by the body, the majority of which is used to produce ATP through glycolysis and mitochondrial oxidative phosphorylation to support neuronal activity and synaptic transmission [31]. We set out to identify the impact of LS2 infection on glucose metabolism in different regions of the brain using ^14^C-labelled 2-DG imaging [22], but instead observed an unprecedented 1.8-fold global *decrease* in glucose utilisation across the whole brain, despite the local presence of glucose-consuming *T. brucei* parasites [12]. Surprisingly, despite the profound changes in brain glucose utilisation and tryptophan levels seen in the S2 murine model, we did not observe any alteration in episodic learning or memory. This is particularly unexpected given the reduced glucose metabolism identified in brain regions including the hippocampus and prefrontal cortex, regions known to contribute to effective NORT performance [32]. Moreover, the lack of an effect on anxiety-like behaviour was surprising given the reduced glucose metabolism seen in brain structures (e.g. amygdala nuclei) involved in this process, and the observation of reduced brain tryptophan, given its established role in anxiety-like behaviour, through its influence on serotonin neurotransmission [33].

Our brain imaging and RNAseq data suggest that HAT infection results in adaptive metabolic reprogramming in the brain, with the cerebral tissue switching away from glucose utilisation to that of using alternative energy sources to maintain brain function. This works to some degree, as reflected by unaltered episodic learning and memory and anxiety-like behaviour during LS2 infection, but impacts on other behavioural measures remain, as evidenced by decreased locomotor activity in our study. Sickness behaviour driven by cytokine release from innate and adaptive immune cells may contribute to the decreased locomotor activity observed during LS2 infection [34].

Bloodstream form *T. brucei* rely on glycolysis to meet their energy demands, with the glycolytic end product pyruvate excreted by the parasite [12]. This pyruvate could be utilised by the host brain tissue as an energy source [35], reducing the host’s reliance on glucose. Whilst transport of pyruvate across the BBB is typically slow it can be increased by disruption of BBB integrity [36], an event which is characteristic of HAT. Interestingly, we found that expression of *Pdk4* was substantially increased in host brain tissue during S1 infection. Pdk4 regulates cellular metabolism by inhibiting the pyruvate dehydrogenase complex (PDC) in mitochondria, slowing down the production of acetyl CoA and ATP through the TCA cycle from pyruvate [37]. Thus, this increased expression may reflect an adaptive response to excess pyruvate availability during S1 infection. Moreover, Pdk4 expression in astrocytes is known to direct pyruvate away from the TCA cycle and supply lactate to brain neurons via metabolic coupling between astrocytes and neurons [38]. This occurs under conditions of stress, when the oxidizing of fatty acids by astrocytes plays a vital role in maintaining neuronal function during periods of high activity. Thus, astrocyte metabolic reprogramming may be initiated during S1 infection to support neuronal function, and this warrants further investigation. Similarly, we saw increased *Hmgcs2* expression during S1 that was reduced during ES2. Hmgs2 is the rate-limiting enzyme for mitochondrial ketogenesis, converting acetyl-CoA and acetoacetyl-CoA into 3-hydroxy-3-methylglutaryl-CoA, which is then processed into ketone bodies (primarily beta-hydroxybutyrate). Upregulated *Hmgcs2* expression may, again, be an adaptive response to increased pyruvate availability from *T. brucei*, as high levels of acetyl-CoA (derived from either fatty acid oxidation or pyruvate metabolism) stimulate Hmgcs2 activity and ketogenesis [39, 40].

Although both *Hmgc2* and *Pdk4* transcript expression returned to naïve levels during ES2 infection, *Ucp2*, which regulates substrate choice for oxidative energy production by mitochondria, decreasing oxidation in glucose-derived pyruvate pathways and enhancing that of fatty acids and glutamine [41], was increased in the brain across the time-course of infection. Thus, the increased expression of *Ucp2* seen in brain tissues during HAT infection may be a key contributor to the decreased cerebral glucose use seen in LS2 animals, biasing mitochondria towards fatty acid metabolism. Interestingly, we also observed an upregulation in *T. brucei* transcripts for glycolysis and pyruvate metabolism in brain-resident parasites in ES2, suggesting that they may capitalise on the host’s reduced use of glucose.

During periods of prolonged fasting, when glucose availability is limited, the brain can utilise ketone bodies produced by β-oxidation of fatty acids as a major alternative energy source [42]. Symptoms of HAT include hypoglycaemia and weight-loss, with depletion of adipose tissue a major contributor [8], and we observed a significant decrease in brain insulin that will result in increased lipolysis [28]. Increased adipocyte lipolysis during HAT has been shown to benefit the host survival [8], and the released fatty acids may contribute to the brain’s energy requirements, reducing the reliance on glucose until fat reserves are depleted. Infection-induced brain inflammation has also been linked to a reduction in brain glucose metabolism that contributes to behavioural changes and delirium [43], characteristic symptoms of S2 HAT. While we did observe decreased locomotor activity in LS2 animals, we found no overt changes in anxiety-like behaviour or in episodic learning and memory in our murine LS2 model. The lack of on impact on the key brain functions could be explained by the metabolism of adipose tissue reserves that were not yet sufficiently depleted to impair brain function, and our humane end points prevented further study of later infections. The alteration of brain glucose metabolism in our LS2 HAT murine model warrants further investigation to understand its clinical significance and its potential link to the adipose tissue loss and neurological perturbance.

In conclusion, we have used a bioluminescent *T. brucei* murine model of HAT to confirm the presence of clinically relevant perturbations in brain tryptophan levels. We have shown that S2 infection leads to adaptive alterations in host brain metabolism, with reduced glucose utilisation and re-programming of mitochondrial metabolism that favours the utilization of fatty acids and lipids to meet the energy demands of the brain, which may also fuel the increased immune activation seen in S2 infection. Despite these significant metabolic changes, host anxiety-like behaviour and episodic learning and memory is not impaired, suggesting that adaptation maintains brain functions and limits the impact of the parasite infection on the brain in S2 infections. These finding may explain the progressive onset of neurological symptoms in HAT patients and drive improvement of the therapeutic outcome for HAT patients who experience persistent neurological symptoms after cure.

## Materials and Methods

### Ethics statement

Mouse studies were carried out under UK Home Office regulations under project license number P15EA559A in compliance with the UK Animals (Scientific Procedures) Act 1986. The work was performed with the approval of Lancaster University Animal Welfare and Ethical Review Body (AWERB) and are reported in line with the Animal Research: Reporting of *in vivo* Experiments (ARRIVE) guidelines. All mice were group housed in individually ventilated cages (Techniplast, UK) on a 12-hour light-dark cycle (lights on at 07:00) at 22 ± 1°C and 65% humidity with access to food and water *ad libitum*.

### Mouse infection and *ex vivo* imaging

Female CD-1 mice (8-12 weeks) were purchased from Charles River (Margate, UK). Two mice were infected intraperitoneally (*i.p.*) with 200 µl phosphate buffered saline (PBS) containing 2 × 10^4^ *T. brucei* VSL2 parasites [19] expressing the red-shifted *Photinus pyralis* luciferase gene (PpyRE9H) [44]. At day 5-post infection, the mice were humanely sacrificed and the infected blood collected, diluted to contain 2 × 10^4^ *T. brucei* cells in 200 µl PBS and injected *i.p.* into each experimental mouse. At specific time-points after the infection (days 7, 14 and 22) mice were injected *i.p.* with 200 µl D-luciferin (Perkin Elmer), and 10 minutes later sacrificed using cervical dislocation and decapitation. Post-mortem tissues including the brain, liver, heart, spleen, muscle, fat, small intestine, colon, eye and mesenteric lymph nodes were removed and placed in 12-well plates. Half of the tissues were used for bioluminescent imaging, and the other half snap frozen in cold isopentane (−40^0^C) and then stored at −80^0^C for metabolomic analysis. Bioluminescent emission from infected tissue was recorded using the ChemiDoc MP Imager (Bio-rad) using Image Lab Software (V6.1, Bio-rad) with an exposure time of 300 s, with a corresponding white light image of the plate captured. Acquired images were processed and quantified using ImageJ by drawing around the tissues and measuring the intensity density [45]. Statistical significance of variance between experimental groups was calculated in GraphPad Prism 10.6.1 using either Welsh’s t-test or a nested 1-way ANOVA with p-Sidaks’ multiple comparison test.

### Tissue sample preparation of targeted metabolomics

Tissues samples from naive and infected animals were defrosted and weighed prior to homogenisation with zirconium beads (BeadBug^™^, Sigma Aldrich) in 1ml of LC-MS grade water in a high-speed mechanical tissue homogenizer (FisherScientific, UK). Homogenised tissues were centrifuged at 15,000 × g at 4^0^C for 10 min, and the resulting supernatant subjected to an acetonitrile/chloroform extraction procedure as described by Lesniak *et al*. [20], lyophilised and stored at −80^0^C until further use.

### High-Pressure Liquid Chromatography (HPLC)

For HPLC analysis an established protocol was followed [20], and samples were analysed using an Agilent Technologies 1260 Infinity II (Böblingen, Germany) HPLC equipped with UV and fluorescence detectors. Samples were run on a TSKgel ODS-80TS column (0.5 µm, 4.6 × 250 mm, from Tosoh Bioscience) with 0.1% TFA in H_2_O: acetonitrile (ACN, 90:10 *v/v*) at a flow rate of 0.8 ml/min and column temperature of 25 ^0^C. Lyophilized samples were dissolved in 1 ml of the mobile phase (0.1% TFA in H_2_0: ACN 90:10 *v/v*). Triplicate technical injections of 100µl for each biological sample were analysed. To compare between control and infected animals, tissues from the same post-infection time-point were run on the same day as one batch to reduce batch to batch variability in relative fluorescence units (RFU). The fluorescence detectors were set at the following channels: for analysis of tryptophan excitation (λ_Ex_) at 297 nm and emission (λ_Em_) at 348 nm, with UV wavelengths set at 220nm, 280nm, 330nm and 364nm. The retention time (RT) for tryptophan of 30.7 minutes was obtained by injecting a reference standard. Data was processed on the BioInert Software (Agilent), then peak integrations were normalised to tissue weight and control group mean. Statistical significance of variance between experimental groups was calculated in GraphPad Prism 10.6.1 using a nested 1-way ANOVA with p-values corrected using Sidaks’ multiple comparison test.

### Liquid Chromatography- Tandem Mass Spectrometry (LC-MS/MS)

A Waters Atlantis T3 (2.1 x 150-mm i.d., 3 µm) reversed-phase column with mobile phases A (0.1% formic acid in water, *v/v*) and B (0.1% formic acid in acetonitrile, *v/v*) was used on the Shimadzu LCMS-8040 instrument. A linear gradient was used for the chromatographic separation of the analytes as previously described by Zhu *et al* [21] with flow rate was 0.4ml/min and column temperature of 35^0^C. The gradient consisted of: 0-10 min from 0% to 40% B; 10-12 min from 40% to 95% B; 12-17 min hold at 95% B; 17-20 min from 95% to 0% B. Analytes were detected in positive ion multiple reaction monitoring (MRM) mode with tryptophan (205.1 > 118.1 *m/z*) eluting with a retention time (RT) of 4.5 min and the internal control 5-methoxy-DL-tryptophan (247.1 > 160.1 *m/z*) eluting with a RT of 5.1 min. For the analysis of samples, lyophilized material was dissolved in 100 µl of mobile phase B (0.1% formic acid in acetonitrile, *v/v*). 10 µg of the internal standard 5-methoxy-DL-tryptophan was added to the recuperated biological samples for each run. Triplicate technical injections of 15 µl for each biological sample were performed. LabSolutions LCMS integrated workstation software (V5, Shimadzu) was used for data collection and peak integration, with integrations normalised to tissue weight and the internal 5-methoxy-DL-tryptophan standard. Statistical analysis was the same as for HPLC. Statistical significance of variance between experimental groups was calculated in GraphPad Prism 10.6.1 using a nested 1-way ANOVA with p-values corrected using Sidaks’ multiple comparison test.

### Open Field Test (OFT)

In preparation for the OFT, CD-1 female mice were placed individually into empty holding cages in order to habituate to the room in which the experiment was held. After 10 min, animals were placed individually into one of the four circular testing arenas, comprising of a black Perspex background (diameter of 38cm, central zone diameter of 12cm) and video recorded for 15 min. Four animals were recorded per run. 70% ethanol was used to clean and remove any prior scent markings before the beginning of the test, and between succeeding runs. A camera was secured to the ceiling and connected to the desktop PC, saving videos for further analysis. The number of animals for each experimental group (infected and control) were day 5 (n = 9), day 12 (n = 10) and day 20 (n = 35).

### Novel Object Recognition Task (NORT)

The day following the OFT, mice were subjected to the Novel Object Recognition Task (NORT). Mice were allowed to habituate to the room in individual holding cages prior to the task. After 10 min, four mice per run were placed individually into the empty arenas and activity recorded for 5 min. Following this, mice were removed and two identical objects (blue pyramids 5.5 × 2.5 × 3 cm, wooden squares 3 × 3 × 3 cm or red oblongs 9 × 3 × 1 cm) were placed into each arena as part of the acquisition phase, placed 15 cm from the arena wall. Mice were allowed 10 min to explore the objects, with their activity being recorded by video, before returning to the home cages. A 24-hour delay was imposed, after which mice were returned to the arenas, now containing one familiar object and a novel object. In this phase, behaviour was video recorded for 10 min.

For both phases of the test, both arenas and objects were cleaned with 70% ethanol between runs. All object allocations to each animal were randomised, as was the side (left or right) of novel object presentation. The number of animals for each group were day 14 (n = 10) and day 22 (n=35).

### Analysis of behavioural videos

Ethovision XT v8.5, Noldus software was used to track and analyse the OFT and NORT videos. For the OFT this allowed parameters such as walking distance, velocity, duration and frequency to be analysed as measures of locomotor activity. Duration and frequency in the centre zone of the arena were analysed as measures of anxiety-like behaviour. The effect of infection and time bin (30 s), on output parameters were analysed using repeated measures Analysis of Variance (ANOVAs) in R software with significance determined at p < 0.05.

### ^14^C-2-Deoxyglucose Functional Brain Imaging

In accordance with previously published protocols by Dawson *et al* [22], cerebral glucose metabolism was determined using ^14^C-2-DG functional brain imaging of naïve (n = 12) and infected mice (n = 22) at day 22. In brief, mice were injected *i.p*. with the 4.635 MBq/kg of the isotope 2-deoxy-D-[^14^C] glucose (dose volume of 2.5 ml/kg) over a 10 second period, diluted in physiological saline. After injection, mice were returned to home cages for 45 min. Subsequently, animals were sacrificed by cervical dislocation followed by decapitation. Brains were dissected out and immediately snap frozen in cold isopentane (−40^0^C) and stored at −80^0^C until further use. A blood sample was taken to measure blood glucose (mmol/L) levels directly using an AccuChek glucose monitor (Aviva).

A terminal blood sample was also collected from each mouse, taken from the neck via torso inversion. Samples were centrifuged at 13,000 rpm to separate the plasma and stored at −80^0^C until required. Liquid scintillation analysis (Packard) was conducted in triplicate to determine plasma ^14^C levels, by placing 10 µl plasma in 1 ml scintillation fluid (Ecoscint XR, National Diagnostics).

Frozen brains were mounted with Shandon^TM^ M-1 Embedding Matrix (ThermoFisher Scientific) and were sectioned coronally at 20 μm within a cryostat (−20^0^C). A series of three consecutive brain slices were collected, thaw mounted onto cover slips (50 × 20mm) and rapidly dried on a hot plate at 70^0^C. The following series of three slices were discarded, and this process was repeated for the whole brain with the exception of the region surrounding the suprachiasmatic nucleus, whereby every section was collected.

The coverslips containing brain slices and ^14^C standards (40-1069 nCi/g tissue equivalents; American Radiolabelled Chemicals, Inc) were opposed to autoradiographic film (Kodak, Biomax MR) within a darkroom for seven days. Then according to manufacturer’s instructions, films were developed using an automated film developer (Konica Minolta, SRX-101A).

### Brain Imaging Analysis

Analysis of the autoradiographic images was conducted using computer-based image analysis (MCID/M5+), following established protocol in Dawson *et al* [22]. Local isotope concentration for each brain region of interest (RoI) was derived from the optical density of the autoradiographic image relative to the co-exposed ^14^C standard. A total of 50 anatomically distinct RoI were chosen (Table S1) and measured with reference to a stereotaxic mouse brain atlas [46]. Each region was measured between 10-12 times per animal (in both hemispheres of 5-6 brain slices). Average whole slice isotope concentration was also measured at each level of the brain analysed, to determine the whole brain average ^14^C level for each animal.

### Tissue Collection and Homogenisation

Brains were dissected and imaged before being placed into RNA Later (Invitrogen) within 10 minutes of mouse death. Brain tissues were left in RNA Later at 4°C for 24 hours then transferred to −80°C for long term storage. RNA was extracted from non-infected controls (n = 6), S1 (day 7, n=6) and ES2 (day 14, n = 4) infected brains and processed in parallel. Brain were defrosted on ice, then excess RNA Later was blotted onto tissue paper before transfer to pre-weighed tubes containing zirconium beads (Benchmark Scientific). Tubes were weighed to record tissue weight, then 1 mL of Trizol reagent was added and the brains were homogenised in a fast prep tissue homogeniser (MP Biomedicals) at 6 m/sec for a 2 min and returned to ice. Samples were centrifuged for 10 minutes at 12,000 × g at 4°C and the fatty layer was removed, then supernatant retained.

### RNA Extraction

Homogenised supernatant was diluted with 100 µL of RNase free water then transferred to Phasemaker tubes (Invitrogen) and incubated at room temperature for 5 min. 200 µL of chloroform was added and the sample shaken vigorously for 15 secs, then incubated at room temperature for 3 min. Tubes were then centrifuged for 5 minutes at 12,000 × g at 4°C, the aqueous phase was transferred to a new tube and an equal volume of 70% ethanol was added and mixed. RNA was extracted using a PureLink RNA Mini Kit (Invitrogen) as per manufacturer’s instructions. RNA quality and quantity was determined using a bioanalyzer (Agilent 2100 Bioanalyzer, RNA 6000 Nano Kit) as per manufacturer’s instructions, with all sample producing an RNA integrity number (RIN) ≥ 9.5. Extracted RNA was stored at −80°C.

### RNAseq Analysis

RNA Sequencing was completed by Novogene using poly-A enrichment prior to strand-specific library preparation and sequencing on an Illumina Novaseq 6000 S4 flow cell with 150 bp end reads providing 6 GB data per sample. Raw reads were pre-processed to remove adaptors, poly-N (>10% N) and low-quality reads (50% < 5 Qscore) using custom Pearl scripts and aligned to the NCBI *Mus musculus* reference genome GRCm39 or the TriTrypDB *Trypanosoma brucei brucei* TREU927 (release 59) genome [47] using HISAT2 (v2.0.5) [48]. The reads counts were calculated using featureCounts (v1.5.0) [49] and converted to Fragments Per Kilobase of transcript sequence per Millions base pairs (FPKM) before differential expression analysis in DESeq2 (v1.20.0) [50]. For a gene to be considered differentially expressed, there must be a greater than 2-fold change in expression (Log_2_ Fold-Change ≥ 1 or ≤ −1) and a calculated adjusted p-value (p-adj) < 0.05. Gene Ontology (GO) enrichment analysis of differentially expressed genes was implemented by clusterProfiler (v3.8.1) [51] for mouse genes or within TriTrypDB for *T.b. brucei* genes [47]. Volcano plots and Principal component analysis (PCA) were generated in Perseus [52].

## Data Availability

RNAseq data has been deposited with the Sequence Read Archive entry PRJNA1366789

## Acknowledgements

The *Trypanosoma brucei brucei* GVR35 strain VSL2 [19] was a gift from Prof. John Kelly (London School of Hygiene and Tropical Medicine).

## Supporting information captions

**Supplementary Figure S1.** Schematic of the murine infection model and study design.

**Supplementary Figure S2.** *Ex vivo* imaging of naïve mouse CD-1 tissues.

**Supplementary Figure S3.** *Ex vivo* imaging of bioluminescent *Trypanosoma brucei* VSL2 in mouse CD-1 tissues at stage I.

**Supplementary Figure S4.** *Ex vivo* imaging of bioluminescent *Trypanosoma brucei* VSL2 in mouse CD-1 tissues at early-stage II.

**Supplementary Figure S5.** *Ex vivo* imaging of bioluminescent *Trypanosoma brucei* VSL2 in mouse CD-1 tissues at late-stage II.

**Supplementary Table S1.** 2-DG function brain imaging data.

**Supplementary Table S2.** RNAseq analysis of mouse brains from naïve and *Trypanosoma brucei brucei* infected mice at stage I and early-stage II.

**Supplementary Table S3** – Mouse Differential Expressed Genes (DEG) in the brain at stage I

**Supplementary Table S4** – Mouse Differential Expressed Genes (DEG) in the brain at early-stage II

**Supplementary Table S5** – RNAseq analysis of *Trypanosoma brucei* within brains of infected mice at stage I and early-stage II

**Supplementary Table S6** – Brain-resident *Trypanosoma brucei* Differential Expressed Genes (DEG) between stage I and early-stage II

## Notes

### Competing Interest Statement

The authors have declared no competing interest.

